# Time Resolved Inspection of Ionizable-Lipid Facilitated Lipid Nanoparticle Disintegration and Cargo Release at an Endosomal Membrane Mimic

**DOI:** 10.1101/2024.02.22.580934

**Authors:** Nima Aliakbarinodehi, Simon Niederkofler, Erik Olsén, Yujia Jing, Gustav Emilsson, Mattias Sjöberg, Björn Agnarsson, Lennart Lindfors, Fredrik Höök

**Affiliations:** Chalmers University of Technology, Department of Physics, Division of Nano and Biophysics, Fysikgränd 3, 41296 Göteborg, Sweden; Advanced Drug Delivery, Pharmaceutical Sciences, R&D, AstraZeneca, 43181 Mölndal, Sweden

**Keywords:** lipid nanoparticle (LNP), mRNA delivery, endosomal escape, endosomal membrane mimic

## Abstract

This study investigates pH-triggered fusion dynamics of lipid nanoparticles (LNPs) with an endosomal membrane mimic, addressing mechanistic aspects of a crucial yet elusive process for effective mRNA delivery. Utilizing time-resolved total internal reflection fluorescence (TIRF) imaging, we observed a delayed onset of LNP fusion upon pH drop, lasting seconds to minutes depending on pH and LNP size. Once initiated, LNP fusion and cargo escape occurred rapidly within tens to hundreds of milliseconds. While LNP disintegration is observed to lead to release of a significant portion of mRNA into the acidic environment, some mRNA molecules remained mobile on the endosomal membrane mimic due to deprotonation-resistant complex salt formation. Comparison of the fusion efficiency of two LNP formulations correlated with protein translation in human primary cell transfection data, emphasizing the importance of biophysical investigations in understanding ionizable-lipid-containing LNP-assisted mRNA delivery mechanisms and providing insights for optimizing mRNA-LNP design for enhanced endosomal escape.

## INTRODUCTION

In mRNA therapeutics, endogenous cellular machineries are utilized to produce therapeutic proteins, thereby providing a promising means to treat a multitude of diseases where conventional medication strategies fail. ^1,2^ To overcome the inherent instability of mRNA and its low capacity to be naturally taken up by cells, a large number of viral and synthetic therapeutic vectors have been developed. ^3–5^ Despite high transfection efficacy, ^6^ viral vectors are associated with challenges related to genetic interference, low cargo payload, toxicity and immunogenicity. ^2,7,8^ This, in turn, has spurred intense efforts in designing nonviral mRNA vectors, as recently manifested by the lipid nanoparticle (LNP) based Covid–19 vaccines developed by Pfizer/BioNTech^9^ and Moderna^10^.

The most efficient LNPs designed for mRNA delivery are formulated using ionizable lipids together with a set of helper lipids, typically cholesterol, gel-phase forming phospholipids, and polyethylene glycol (PEG) modified lipids. ^11^ Efficient mRNA encapsulation, appropriate LNP structure and desired stability^12,13^ is typically obtained by tuning the ratio between mRNA and the lipid components utilizing microfluidic assisted rapid mixing precipitation protocols. ^14^ This approach has been demonstrated to generate LNPs that display successful endocytic uptake accompanied by mRNA-assisted protein expression, low clearance and degradation and even specific cell targeting have been demonstrated. ^4,15^ Yet, LNPs offer significantly lower transfection efficacy than their viral counterparts, ^6^ being attributed to both extra- and intracellular obstacles, of which the endosomal escape event has been identified as a key bottle neck. ^16,17^ This process, during which mRNA is translocated across the endosomal membrane into the cytosol of the target cell, depends on the gradual acidification of the endosomal environment,^18,19^ which in turn is believed to promote electrostatic attraction between the cationic ionizable lipid contained in LNPs and the anionic endosomal membranes, eventually resulting in LNP disintegration and subsequent mRNA translocation across the endosomal membrane. However, even if cellular endocytic LNP uptake is usually very efficient, functional mRNA delivery is not; in fact, with ionizable-lipid containing LNPs designed for siRNA delivery, less than 2% of the endocytosed cargo resulted in a functional response^11,20^, and the efficacy is even lower in the case of high molecular-weight mRNA.^21^

Insights of this type are typically obtained using advanced optical imaging approaches utilizing *in vitro* cellular assays, ^6,17,21,22^ which are further corroborated through *in vivo* studies. ^23^ Recent work have shown that fundamental mechanistic insights with respect to the nature of LNP interactions with cellular membranes can also be gained by making use of simplified mimics of the anionic endosomal membrane. For example, pH-induced binding of LNPs to anionic lipid monolayers formed at an air-water interface revealed that pH-induced lipid transfer induces structural alterations of endosomal membrane mimics.^24^ By forming a supported lipid bilayer (SLB) containing 6 mol% POPS on a planar glass substrate, thus mimicking the anionic character of the endosomal membrane, it was shown that pH-induced LNP binding to the membrane is accompanied by ionizable-lipid transfer that leads to charge-neutralization of the endosomal membrane, presumably relevant in the context of endosomal arrest. ^25^

In this work, we have used time-resolved dual-color total internal reflection fluorescence (TIRF) microscopy to investigate pH-induced interactions between individual LNPs and a supported endosomal membrane mimic. Inspired by previous work demonstrating that investigations of membrane-enveloped-virus fusion benefit from minimizing the contact between the SLB and the underlying support, ^26,27^ the endosomal membrane mimic was formed on a porous silica substrate, ^28,29^ previously shown to display significantly higher lipid diffusivity than when formed on planar glass, and also to be compatible with lipid molecular translocation across the lipid membrane. ^30^ Since cellular endocytotic LNP uptake is believed to be mediated by the specific binding between ApoE spontaneously adsorbed on the LNP surface in the presence of serum proteins and LDL receptors present on the surface of recipient cells, ^31,32^ LNPs are expected to reside in close proximity to the endosomal membrane. The LNPs were therefore molecularly anchored to the endosomal membrane mimic using a NeutrAvidin-biotin linkage, which also enabled continuous time-resolved imaging of LNPs during the gradual endosomal acidification process to be simulated by varying the pH of the bulk solution using microfluidic assisted liquid exchange.

The investigation was primarily focused on a particular LNP formulation containing DLin-MC3-DMA as the ionizable lipid, and cholesterol, DSPC and DMPE-PEG2000 serving as helper lipids, which in previous *in vitro* cellular assays was demonstrated to display efficient cellular uptake and high protein expression levels. ^33^ The LNP fusion kinetics and the fate of the LNP cargo was microscopically visualized by staining the LNPs with 0.06 mol% Rhodamine-labeled DOPE (Rhod-DOPE) and with 20% of the mRNA being Cy5 labeled (Cy5-mRNA). Individual fusion events were statistically analyzed with respect to i) Rhod-DOPE and Cy5-mRNA release kinetics, ii) the pH-dependency of the diffusivity of individual mRNA and mRNA clusters that remained attached to the endosomal membrane mimic after completed LNP fusion, and iii) the wait time observed between the rapid (< 1 second) pH drop and the actual onset of LNP fusion with the endosomal membrane mimic.

The LNP fusion efficiency was also compared to an additional LNP formulation, designed to display a two times higher surface concentrations of the gel-phase forming DSPC lipid, a variation previously shown to display similar cellular uptake efficiency, but more than one order of magnitude lower protein expression levels.^33^ The mechanistic insights related to pH-induced LNP disintegration revealed through this investigation are discussed in the context of possible endosomal escape pathways and how the use of simplified biomimetic assays may help advancing the design of more efficient mRNA delivery formulations.

## RESULTS

### The pH dependence of the LNP fusion efficiency

The anionic endosomal membrane mimic was formed on nanoporous silica with pore dimensions of around 6 nm through lipid vesicles adsorption-induced SLB formation^28^ using lipid vesicles composed of 93.5 mol% POPC, 6 mol% POPS, representing the negative charge of the early to late endosomal membrane. ^34^ To enable visualization of the SLB formation process using TIRF microscopy and to perform lateral mobility determinations using fluorescence recovery after photobleaching (FRAP), ^35^ the lipid vesicles contained 0.5 mol% NBD-labeled lipids (NBD-DOPE). The TIRF and FRAP analysis revealed successful formation of a continuous SLB with a diffusivity constant *D* of 4.4 ± 0.3 µm^2^s^-1^ (n >3) and an immobile fraction of 0.06 ± 0.02 (n >3). This is at least twice the diffusivity typically obtained for SLBs with the same lipid composition formed on planar glass, ^25^ attributed to reduced lipid pinning at the interface between the SLB and the nanoporous silica support.

To enable time-resolved TIRF imaging of individual LNP fusion events, LNPs with a number average diameter and polydispersity index (PDI) of 140 nm and < 0.1, respectively, were formulated using 53.47 mol% ionizable cationic lipids (DLin-MC3-DMA), 4.65 mol% DSPC, 41.114 mol% cholesterol, 0.7 mol% PEG-modified lipids (DMPE-PEG2000), 0.06 mol% fluorescent Rhodamine labeled lipids (Rhod-DOPE), and 0.006 mol% biotin-modified lipids (DSPE-PEG2000-Biotin), while the eGFP-encoding mRNA cargo was composed of Cy5-labeled mRNA (Cy5-mRNA) and non-labeled mRNA at a 1:4 ratio (Methods and Supporting Information, Figure S1 and Table S1). The LNPs were bound to the endosomal membrane mimic using NeutrAvidin as a linker between biotin-PEG-lipids in the LNPs (∼70 per LNP) and 0.05 mol% of Biotin-Cap-DOPE in the SLB, as schematically depicted in Figure 1a.

Binding of biotin-modified LNPs to the NeutrAvidin-modified endosomal membrane mimic formed on nanoporous silica (Figure 1b) was recorded using dual color TIRF microscopy at 3 frames per second (fps) upon LNP injection at a concentration of ∼10^9^ LNPs/mL in a flow cell (3.8×17×0.4 mm in width × length × height) at a volumetric flow rate of 140 µL min^-1^ until an LNP coverage of ∼0.03 μm^-2^ (∼600 LNPs per field of view) was reached, typically within 5 to 10 minutes. After terminating the LNP binding by rinsing the channel with a buffer solution at pH 7.5, the pH of the flowing solution was subsequently changed via rapid liquid exchange (<1 second within the field of view) from 7.5 to 6.6, 6.0, 5.6, while continuously recording LNP fluorescence emission for around 5 minutes at each pH value.

The most dramatic response was observed upon reducing the pH from 6.6 to 6.0, resulting in a drastic decrease in the number of LNPs with a detectable Rhod-DOPE emission signal, as illustrated in Figures 1c, d. A cumulative sum of all individual fusion events versus time upon the subsequent pH reduction steps (Figure 1e) shows that very few fusion events are detected when the pH is reduced from 7.5 to 6.6, while the majority of fusion events occur after a wait-time of ∼10s and within less than 100 seconds upon reducing the pH from 6.6 to 6.0. Inspection of the micrographs revealed fusion efficiencies relative to the total number of bound LNPs at pH 7.5 of around 3, 54 and 10% at pH 6.6, 6.0 and 5.6, respectively (Figure 1f).

To estimate the degree of ionization of the LNPs, that is, the process that is responsible for inducing electrostatic attraction to the anionic membrane, the fraction of ionized Dlin-MC3-DMA as a function of pH was measured using anionic fluorescent dye 2-(p-toluidino)-6-napthalene sulfonic acid (TNS), which undergoes a significant fluorescent enhancement when binding to positively charged lipids.^36,37^ The TNS assay displays a relatively sharp transition around an inflection point at pH 6.6, with 20 and 80% of Dlin-MC3-DMA available for TNS binding being ionized at around pH 7.5 and 6.0, respectively (Figure S2). While it remains uncertain whether non-surface-exposed ionized Dlin-MC3-DMA is available for TNS binding, these results suggest that substantial ionization of Dlin-MC3-DMA is required to initiate electrostatically driven fusion between ionized LNPs and the anionic endosomal membrane mimic.

**Figure 1.**
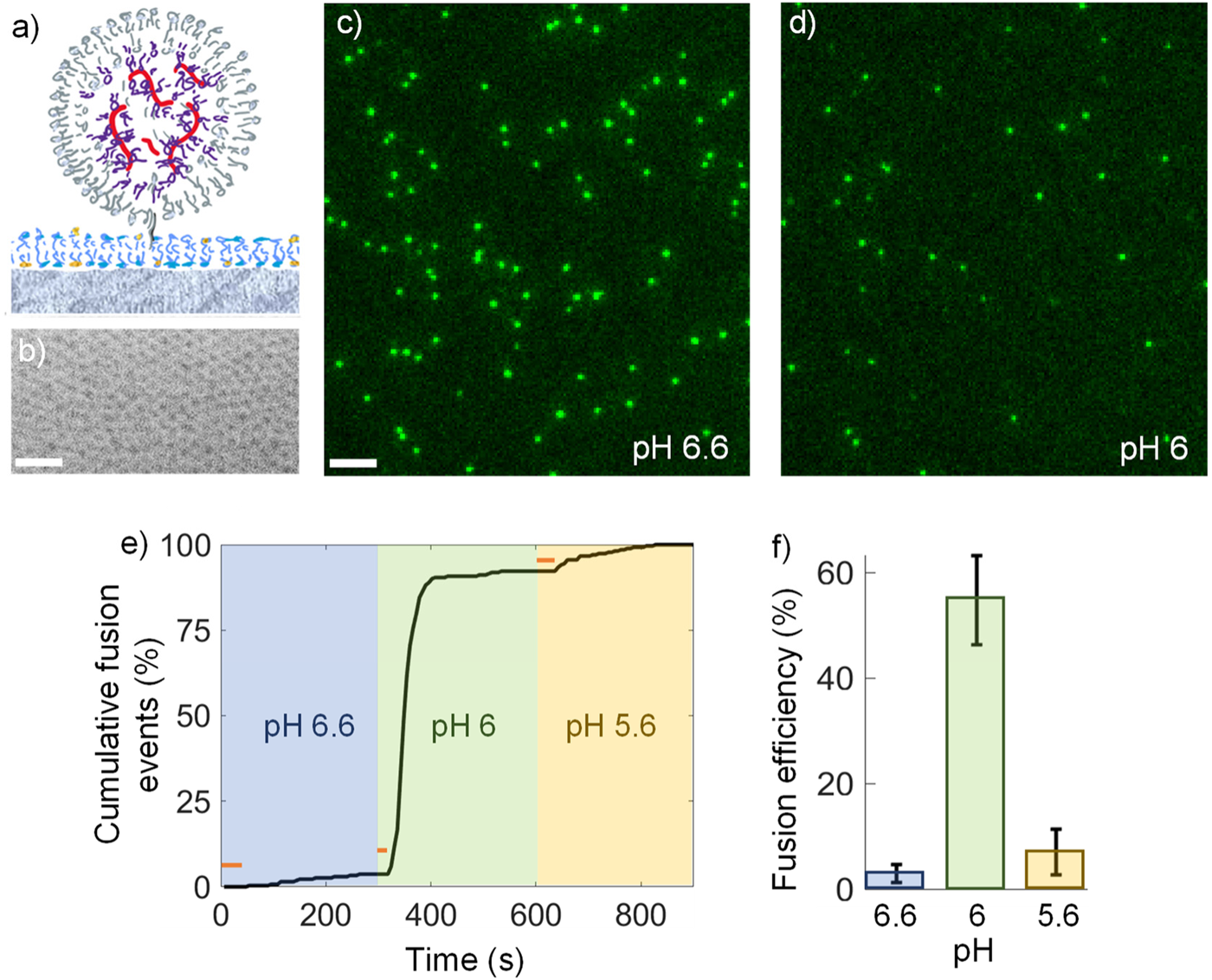
**a)** Schematic illustration of a biotin-modified LNP bound via Neutravidin-linking to a planar biotin-lipid containing endosomal membrane mimic formed on a nanoporous silica substrate. **b)** SEM image showing the estimated 6 nm pore size of the porous silica substrate (scale bar 50 nm). **c)** and **d)** TIRF micrographs of tethered LNPs (Rhod-DOPE emission) 5 minutes after exposure to c) pH 6.6 (scale bar 5 μm) and d) pH 6.0 (scale bar 5 μm). **e)** Cumulative number of fusion events displayed in percentage of total fusion events versus time upon subsequent reductions in pH from 7.5 to 6.6, 6.0 and 5.6. The red lines indicate the wait-time between the reduction in pH to the first recorded fusion event. **f)** Fusion efficiency versus pH displayed in relation to the total number of initially tethered LNPs.

### The spatiotemporal dynamics differ between lipids and cargo upon LNP fusion

The temporal evolution of the LNP fluorescence emission signal upon the actual fusion event is characterized by a synchronized reduction in both Rhod-DOPE and Cy5-mRNA emission signals (lower LNP in Figures 2a, b). However, the amount of signal-reduction differs significantly, and from analyzing a two-dimensional Gaussian function fitted to the background-subtracted emission-profile of individual LNPs (Figures 2c, d), as previously described for the analysis of single lipid-vesicle fusion events^27^, significant differences in the spatio-temporal evolutions for the two fluorescent signals are revealed (Figures 2e, f).

Considering the Rhod-DOPE emission signal first, it displays a gradual reduction at a time scale on the order of a few hundred milliseconds (green curve in Figure 2e). This decrease is accompanied by concurrent increase in the Rhod-DOPE emission intensity in the area surrounding the docking site of the LNP (dashed curve in Figure 2e). The time evolution of the variance, σ^2^, of the fitted Gaussian function reveals a mean diffusion coefficient *D* for Rhod-DOPE of 2.6 ± 0.9 µm^2^ s^-1^ (Figure 2g), and a reduction in the total emission intensity by more than 90% (Figure 2h). These findings are indicative of near complete escape of Rhod-DOPE into the endosomal membrane mimic, in analogy with pH-induced fusion of enveloped viruses to membrane mimics when visualized using dye-labeled lipids.^26^ The somewhat lower diffusivity of Rhod-DOPE compared to the unperturbed endosomal membrane mimic of ∼4.4 µm^2^ s^-1^ is attributed to the nature of the lipid-constituents of the LNP, including both DLin-MC3-DMA, DSPC and cholesterol, the latter of which is known to reduce membrane mobility.

In contrast, the Cy5-mRNA emission signal displays a rather abrupt drop (< 300 ms) upon fusion, resulting in an intensity reduction of around 70% (red curve in Figure 2h), but unlike the dye-labeled Rhod-DOPE lipid, the Cy5-mRNA signal reveals no detectable signs of a concurrent increase in the area around the docking site of the LNP (dashed line in Figure 2f). This suggests that the measured response is caused either by i) mRNA translocation across the endosomal membrane-mimic-interface into the porous regions of the substrate, where the illumination intensity is approximately 4 times lower than the TIR illumination intensity at the nanoporous interface,^38^ or ii) photophysical changes of the Cy5 emission induced upon LNP collapse, or iii) Cy5-mRNA release into the acidic bulk solution above the endosomal membrane mimic, ^39^ or a combination of all these processes.

Inspection of fusion kinetics employing epi illumination, which ensures a consistent illumination across the nanoporous interface, revealed similar kinetics (Figure S3) to that shown in Figure 2f, with no apparent indication of Cy5-mRNA being translocated across the membrane and into the nanoporous substrate, suggesting that this process is most likely not the predominant contribution to the observed reduction in fluorescence intensity. Considering photophysical changes, it should be noted that each Cy5-mRNA contains on average ∼34 Cy5 dyes per mRNA (see Materials and Methods). With 20% of the mRNA cargo being labeled, and with on average 200 mRNA per 140 nm diameter LNP,^33^ the mean distance between Cy5 dyes within an LNP becomes ∼10 nm, which is significantly larger than the 6 nm Förster distance of Cy5.^40^ Thus, even if the LNP fusion would lead to complete expulsion of its internal 25% volume of water,^33^ the accompanied reduction in intermolecular distance between adjacent Cy5 molecules will cause a decrease in fluorescence emission intensity due to quenching of 5 to 10% at most. We thus exclude also photophysical effects as the primary cause of the observed reduction in Cy5-mRNA emission, leaving mRNA release into the solution above the endosomal membrane mimic as the most plausible cause for the rapid and dramatic decrease in the Cy5 signal intensity.

The remaining fraction of the Cy5-mRNA intensity (∼30%) continued to reside at or near the LNP docking site after fusion, which can presumably be attributed to entangled mRNA being electrostatically bound to the positively charged head group of DLin-MC3-DMA. This molecular complex may in turn experience hydrophobic association with lipid assemblies at the site of collapse, which is likely to restrict mRNA translocation across the membrane. However, it is important to keep in mind that the reported dye-labeled components only represent a small fraction of the molecules participating in the fusion process, which adds uncertainty to any presumptions made to the behavior of the unlabeled components. One way of addressing this issue is to apply dual-color-fluorescence and label-free-scattering microscopy to study the correlation in scattering and fluorescence intensity of the LNPs upon pH changes.^41,42^ Although this method is currently limited to investigations on planar glass substrates, where the LNP fusion efficiency with the endosomal membrane mimic is observed to be an order of magnitude lower (<10%) than that reported here on porous silica substrates, a similar reduction in Rhod-DOPE emission was observed upon pH-induced fusion of LNPs tethered to the same type of supported endosomal membrane mimic (Figure S4). Further, it was evident from changes in the label-free scattering signal that an LNP fusion event is accompanied by a reduction in the scattering intensity by more than 90% (Figure S4). Since the scattering intensity is to a first approximation proportional to the square of the LNP mass,^42^ this indicates that a large fraction of the LNP content (∼70% of the mass) dissolves into the endosomal membrane mimic and/or escapes into solution upon fusion.

The analysis depicted so far suggests that at least the majority of the lipid material of the LNPs is efficiently integrated into the underlying endosomal membrane mimic during pH-induced LNP fusion. To specifically trace the fate of DLin-MC3-DMA during LNP fusion and disintegration, experiments were conducted using LNPs formed without Rhod-DOPE, but instead with calcein, a highly negatively charged dye^43^ that is expected to be electrostatically bound to the ionized headgroup of DLin-MC3-DMA lipid upon LNP formation at low pH. Even though the nucleotide (PolyA) encapsulation efficiency was in this case reduced by a factor of 3 (Table S1), the release kinetics of calcein observed upon pH induced LNP fusion (Figure S5) was similar to that observed for Rhod-DOPE, suggesting that a significant fraction of the calcein is transferred into the endosomal membrane in the form of calcein-DLin-MC3-DMA complexes, which is further supported by an average diffusion constant of 2.3 μm^2^s^-1^, being similar to that obtained for Rhod-DOPE (Figure 2g).

**Figure 2.**
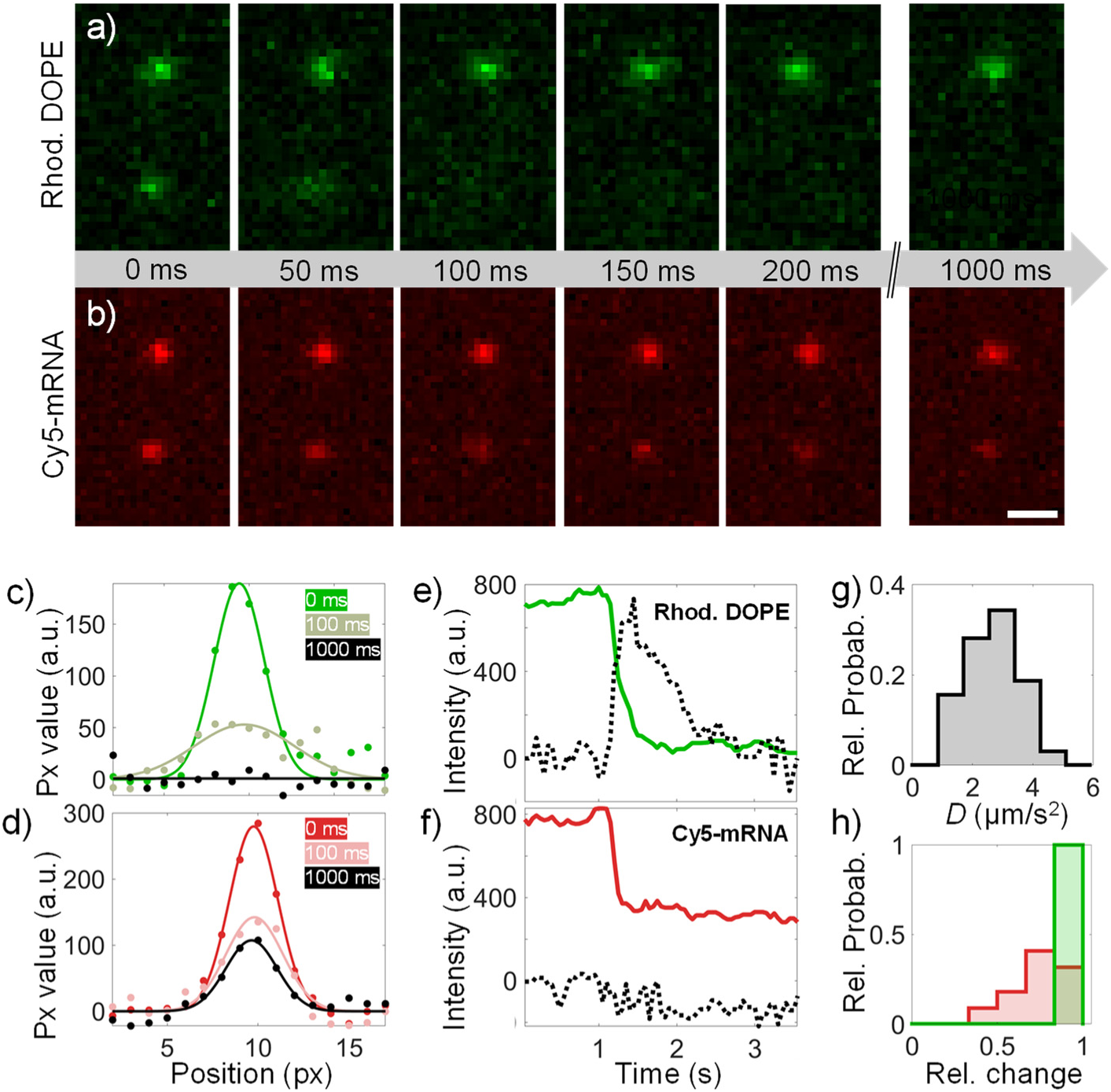
TIRF micrographs showing **a)** Rhod-DOPE and **b)** Cy5-mRNA emission signals for two tethered LNPs measured at 20 fps upon a reduction in pH from 7.5 to 5.6 (scale bar 1 μm). The upper LNP does not show any respond to the pH change in this time interval while the lower does. Time zero in a) and b) is defined as the time of onset of pH change. Background-subtracted spatio-temporal emission profiles for individual LNPs were fitted to two-dimensional Gaussian profiles that are in **c)** and **d)** represented as one-dimensional averages for Rhod-DOPE and Cy5-mRNA, respectively. These were further used to display the temporal evolution of the total LNP emission intensity (solid lines) and corresponding emission from and area surrounding the LNP docking site (dashed lines) for **e)** Rhod-DOPE and **f)** Cy5-mRNA. **g)** Diffusion constants, *D,* obtained from the variance, σ^2^ = 2*Dt,* of the time evolution of the two-dimensional Gaussian emission profiles from 32 LNPs.[40] **h)** Relative change 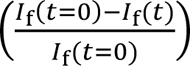 in the Rhod-DOPE (green) and Cy5-mRNA (red) emission intensities for LNPs that undergo pH-induced fusion.

### mRNA remains bound to the endosomal membrane mimic after LNP fusion

Considering molecular transfer from lipid-based drug carriers during interactions with endosomal membrane mimics, it has been previously reported that siRNAs escape into bulk upon pH-induced electrostatic interactions with charged liposomes designed to mimic the anionic character of the endosomal membrane, ^44^ and individual ssDNAs formulated in DOTAP-ssDNA lipoplexes have been shown to remain electrostatically bound to the cationic DOTAP lipids upon electrostatically driven fusion between DOTAP-ssDNA lipoplexes and anionic SLBs.^45^ It is therefore not unlikely that mRNA form similar complexes with DLin-MC3-DMA lipids, which, if escape into solution occurs, could subsequently rebind to the supported endosomal membrane via hydrophobically driven insertion. Alternatively, suspended mRNA could rebind via electrostatic association with DLin-MC3-DMA that has escaped into the endosomal membrane mimic. For the latter scenario to be plausible, rebinding is expected to occur near the LNP docking site soon after fusion, before the positive charge of DLin-MC3-DMA is counter-balanced by negatively charged POPS lipids in the endosomal membrane mimic. To further elucidate the fate of Cy5-mRNA upon pH induced fusion, identical set of experiments were conducted, employing high-intensity epi illumination at an acquisition rate of 2 fps, facilitating the detection and tracking of weak fluorescence signals. To minimize bleaching effects, the measurements were initiated 5 minutes after subsequently induced pH reductions, that is, when essentially no more fusion events were observed (see Figure 1e).

The results are summarized, together with representative micrographs (Figure 3a), as scatter plots of the Cy5-mRNA emission intensity plotted versus lateral diffusivity for all individual detectable entities, recorded after initial LNP binding at pH 7.5 (Figures 3b), and at pH 6.6, 6.0, and 5.6 (Figures 3c to e, respectively), followed by an exchange back to pH 7.5 (Figure 3f). After initial LNP binding at pH 7.5, the majority of detected entities (∼97%) display a rather uniform intensity distribution with a half width maximum value corresponding to around 50% of the peak value, which is typically observed for individual LNPs of this type, ^46^ but also a small fraction (3%) of entities with around 1.5 times lower intensities when displayed on log-log scale (Figure 3b). With around 40 Cy5-mRNA molecules per LNP (see above), an individual Cy5-mRNA is expected to have around 1.6 times lower intensity when plotted on a logarithmic scale, suggesting that that the low intensity detections are dominated by individual membrane bound Cy5-mRNA.

Upon sequential reduction in pH from 7.5 to 5.6 (Figures 3b-e), there is a gradual shift from the high intensity distribution towards lower intensities, with the most significant change occurring when the pH is lowered from 6.6 to 6.0 (Figure 3d). In this step, there is also a significant increase in the number of individual Cy5-mRNA detections, increasing from 11% to 40% of the detected entities. These observations are attributed to the fact that the fusion efficiency is highest in this step (Figure 1), resulting in a substantial fraction of mRNA being released into solution; however, as evident from this analysis, a small fraction of individual mRNA also rebinds to the endosomal membrane mimic. Also note that there is a reduction in the number of individual Cy5-mRNA detection events at pH 5.6 compared with 6.0 (Figure 3e), tentatively attributed to a reduction in negative charge of POPS near pH 5.^47^

Considering the estimated diffusion constants, we refrain from attempts to quantify the diffusivity in terms of number of contact points between the mobile entities and the endosomal membrane mimic since the diffusivity of the endosomal membrane mimic may change in response to lipids escaping from fusing LNPs. Note, though, that the diffusion constants of individual mRNA ranges from values similar to those measured for individual lipids (1 to 4 μm^2^s^-1^) to orders of magnitude lower values, being significative of multiple contact points with the endosomal membrane mimic. Yet, insights can be gained from inspecting relative changes in the diffusivity distributions when the pH is sequentially reduced from 7.5 to 5.6. First, the diffusivity of the high-intensity distribution shifts towards significantly lower values already when the pH is reduced from 7.5 to 6.6, which can be attributed to an increase in electrostatic attraction between partially ionized but non-fused LNPs and the anionic endosomal membrane. In contrast, there are no dramatic variations in diffusivity distribution of individual Cy5-mRNA, except for a reduction in the fraction of mRNA with high diffusivity at pH 5.6, which is attributed to mRNA release due to a reduction in the electrostatic attraction between mRNA and POPS.

These observations might indeed be relevant in the context of endosomal escape, since it suggests that mRNA could in fact be associated with the endosomal membrane when being translocated to the neutral cytosolic environment. To simulate this step, that is, mRNA translocation from an acidic to a neutral environment, the pH was finally increased from 5.6 to 7.5. Intuitively one would expect that this would lead to complete deprotonation of DLin-MC3-DMA accompanied by release of Cy5-mRNA into the bulk solution. Instead, although the diffusivity of individual Cy5-mRNA increased (Figures 3f), signaling partial decoupling from the membrane, the fraction of single Cy5-mRNA increased from 26 to 42%, accompanied by a reduction of the intensity distribution of the high intensity population (Figure 3f). This observation indicates release of Cy5-mRNA from the site of LNP fusion which, at least in part, remains membrane bound as Cy5-mRNA monomers. This phenomenon is in turn attributed to complex salt formation between mRNA and Dlin-MC3-DMA, which was recently shown to remain stable in LNPs, also at neutral pH.^48^

**Figure 3.**
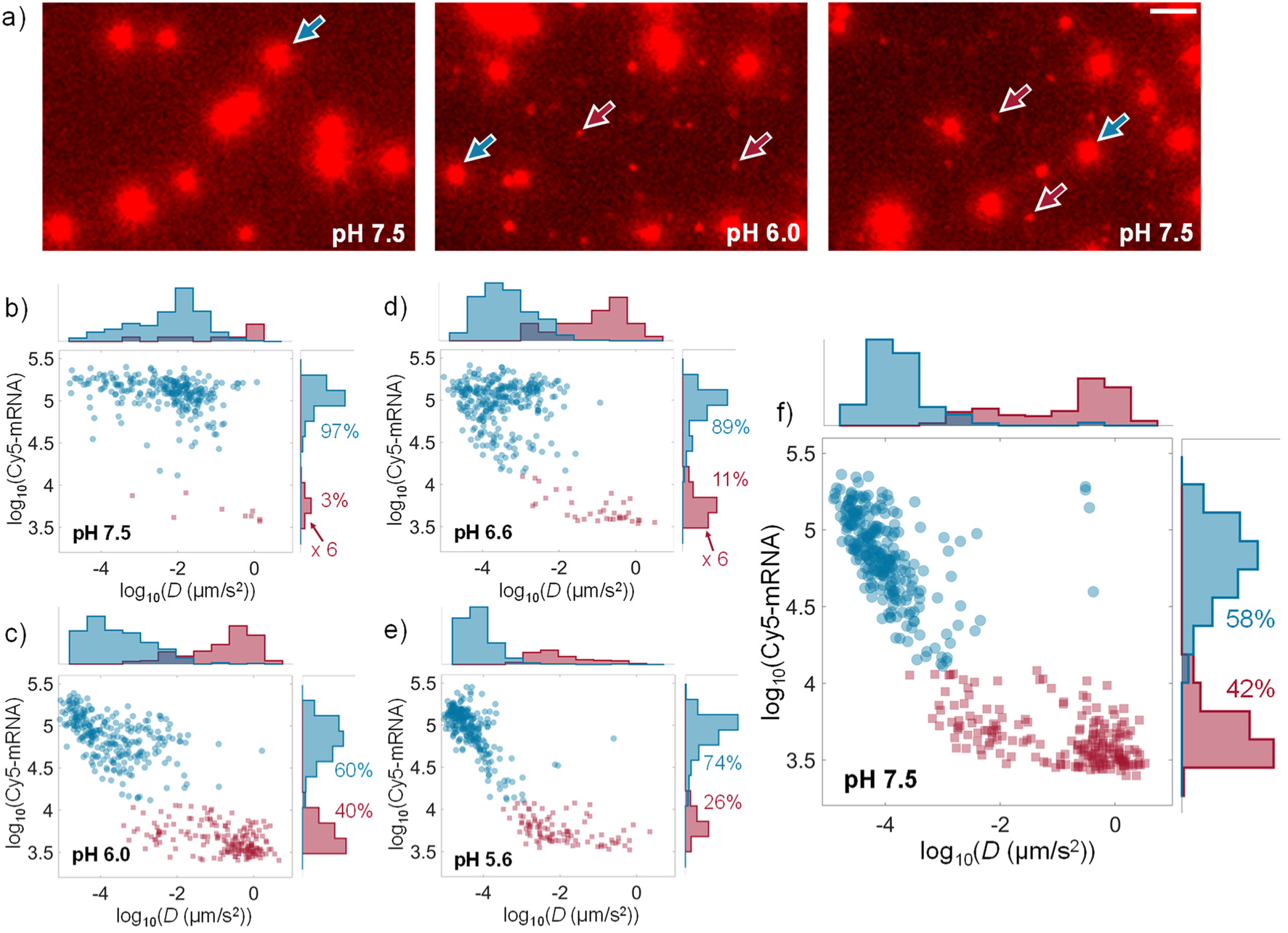
**a)** Cy5-mRNA epi fluorescence micrographs (from different fields of view) displaying examples of individual LNPs with high intensity and low mobility (indicated with blue arrow) and low intensity and high mobility (indicated by red arrows) and upon tethering at pH 7.5 (top), after reduction to pH 6.0 (middle) and after reverting back to pH 7.5 (bottom). The scale bar corresponds to 5 μm. b-f) Log-log representation of Cy5-mRNA emission intensity versus diffusion constant *D* for individual detections at pH **b)** 7.5, **c)** 6.6, **d)** 6.0, **e)** 5.6 and **f)** after reverting pH back to pH 7.5. To guide the eye the data is color-coded based on fluorescence intensity.

### The pH dependence of LNP fusion depends on LNP formulation

The investigation presented above focuses on an LNP formulation that was previously shown to induce efficient mRNA transfection efficiency.^33^ This previous work also showed that by approximately doubling the fraction of the gel-phase forming DSPC lipid, known to be predominantly located at the surface of this particular type of LNP, the protein production was reduced by more than one order of magnitude. Since the cellular uptake of these two types of LNPs was observed to not differ significantly, the difference in protein production was attributed to a reduction in endosomal escape efficiency. Since the endosomal escape event is likely to be closely connected with the capacity of LNPs to fuse with the endosomal membrane in response to a reduction in pH, these results motivated us to explore if the minimalistic LNP fusion assay presented in this work could also help elucidating differences in the fusogenicity of these two types of LNPs (see Materials and Methods and Table S1).

While neither of the two LNP formulations display significant fusion when the pH was dropped from 7.5 to 6.6 (< 5%), the LNPs containing low DSPC concentration (low-DSPC LNPs) displayed at least three times higher (∼55%) fusion efficiency than the high-DSPC LNPs (∼17%) upon a reduction of the pH from 6.6 to 6.0, with cumulative fusion efficiencies of ∼62 and 21%, respectively, upon subsequent reduction of the pH to 5.6 (Figure 4a). It is also worth noting that low- and high-DSPC LNPs contain 0.7 and 0.25% DMPE-PEG(2000), respectively, which converts to distances between surface associated PEG chains of approximately 2.5 and 4.1 nm, respectively. With a Flory radius of ∼3.5 nm for 2 kDa PEG,^49^ this suggests a PEG brush conformation for low-DSPC LNPs, which should intuitively prevent close contact with the endosomal membrane mimic. Yet, under the reasonable assumption that DMPE-PEG2000 remains bound to the LNPs after surface attachment, the results shows that potential steric repulsion induced by the presence of PEG seems to be overcome by the pH-induced electrostatic attraction, and that high surface concentration of the gel-phase forming DSPC lipid is the dominating reason for the lower fusion efficiency of high-DSPC LNPs. This observation also supports that the previously reported^33^ difference in protein production for these two LNPs is indeed most likely due to a difference in endosomal escape efficiency caused by the difference in surface concentration of DSPC.

Further, by exploring differences in the fusion behavior at different pH, additional mechanistic insights can be gained. Figures 4b and d summarize the key results of this set of experiments in scatter plots displaying the Rhod-DOPE fluorescence emission, which to a good approximation represents a measure of LNP size,^46^ prior to fusion at pH 7.5 versus the wait time (Figure 1e) between the rapid pH drop (< 1 second) and the onset of fusion, together with histograms projected towards the respective axis. Focusing on the low-DSPC LNPs first (Figure 4b), the average wait time is on the order of 200 seconds upon reducing of the pH from 7.5 to 6.6, which is around 10 times longer than the corresponding average wait time for the fusion events observed upon subsequent reduction of the pH from 6.6 to 6.0 (around 20 seconds). It is also clear that upon reducing the pH from 7.5. to 6.6, it is preferentially small (low Rhod-DOPE emission) LNPs that undergo fusion, while at pH 6.0, the LNPs that undergo fusion display a size distribution that is similar to that of the original distribution at pH 7.5. In contrast, for high-DSPC LNPs, the fusion efficiency is markedly lower and the statistics therefore low, but it is still clear that there is a broad distribution of wait-times at all pH, with a tendency towards increased fusion efficiency and shorter wait-times for small (low Rhod-DOPE emission) LNPs (Figure 4c).

Despite these differences between low- and high-DSPC LNPs, the TNS assay shows essentially identical transitions around an inflection point at pH 6.6 for both types of LNPs. (Figure S2). Under the assumption that DLin-MC3-DMA is homogeneously distributed within the LNP at neutral pH and that the fraction of ionized DLin-MC3-DMA is independent of LNP size, the surface coverage of ionized DLin-MC3-DMA that can potentially be reached should scale as the inverse of LNP radius. One plausible explanation to the observation showing that small LNPs tend to fuse more efficiently at pH 6.6 (Figures 4b, c), at which only a fraction of DLin-MC3-DMA is ionized, is therefore that only LNPs in the smaller size regime gain sufficient surface charge for the electrostatic attraction to become high enough for fusion to occur. However, if one assumes that up to one third of DLin-MC3-DMA can be engaged in a complex salt with mRNA,^48^ there is on the order of 450×10^3^ DLin-MC3-DMA available for ionization within a 140 nm diameter LNP, while the maximum number of lipids (assuming a head group area of ∼1 nm^2^) located at the LNP interface is around 60×10^3^. Thus, if 50% of the DLin-MC3-DMA is ionized already at pH 6.6, the amount of ionized DLin-MC3-DMA is unlikely to be the limiting factor for fusion. Rather, the fusion event seems to be limited by the capacity of DLin-MC3-DMA to be sufficiently and favorably exposed at the LNP interface, which is likely to include a process in which both DSPC, cholesterol and PEGylated lipids are replaced or expelled. Such mechanisms may very well vary across LNP size, and also be influenced by size-dependent variations in the surface concentration of gel-phase forming DSPC as well as differences in interfacial membrane strain.

It is also worth noting that although the fusion efficiency of low-DSPC LNPs is as high as 60%, there is still a significant fraction of the tethered LNPs that does not undergo fusion. Together with the variations in wait-time and loss of mRNA observed between individual low-DSPC LNPs, this suggests that not only are there significant differences between the two types of LNPs investigated in this work, but also a significant heterogeneity within an individual LNP population. This is indeed evident also for the small fraction of high-DSPC LNPs that undergo fusion, which display significant variations with respect to the wait-time prior to fusion; yet, irrespective of LNP type and size, essentially no fusion events were observed with wait-times shorter than around 5 to 10 seconds even at pH 6.0 and 5.6 (Figures 4b, c), suggesting that also for LNPs with “optimal” structure it takes several seconds for ionized DLin-MC3-DMA to be favorably exposed on the LNP interface for fusion to occur.

**Figure 4.**
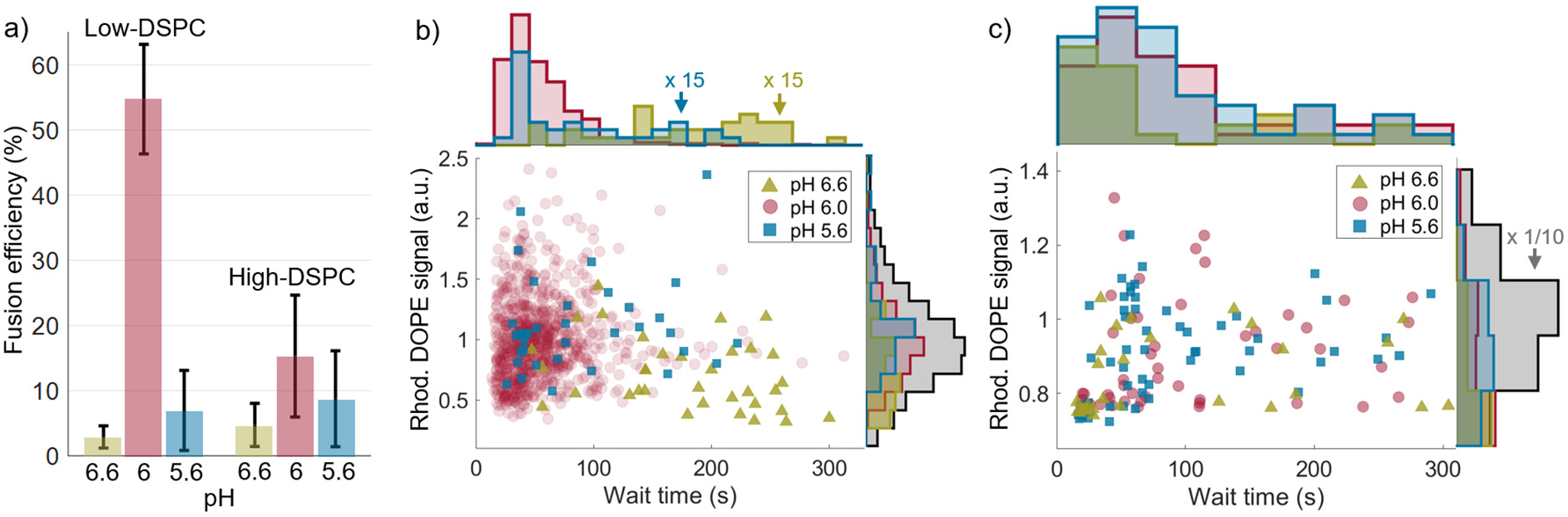
**a)** Fusion efficiencies at different pH for low-DSPC- and high-DSPC LNPs with error bars representing standard deviation from three measurements. Scatter plots displaying the Rhod-DOPE fluorescence emission intensity at pH 7.5, prior to fusion, versus the waiting time to fusion onset together with histograms projected on the respective axis for **b)** low-DSPC LNPs and **c)** high-DSPC LNPs. Data extracted from 3 independent measurements.

## DISCUSSION

The minimalistic model of the anionic endosomal membrane that was in this work designed and used to investigate the temporal dynamics of pH-induced LNP fusion with, and subsequent disintegration at, an endosomal membrane mimic demonstrates common general features, but also significant differences between different types of LNPs as well as heterogeneity between individual LNPs of the same type. Given the simplicity and homogeneity of the endosomal membrane mimic used here compared with a natural endosomal membrane, which contains a much more complex lipid composition as well as a many different types of membrane proteins,^50^ it is not unlikely that the heterogeneity with respect to LNP fusion is even higher in a natural cellular environment. This holds especially true since LNPs also become decorated with a protein corona prior to endocytic uptake,^51^a process that is known to influence their pH-dependent interaction with endosomal membrane mimics similar to the one used in this work. ^25^ Yet, the fundamental mechanisms and distinct features observed in this work are likely to resemble those occurring in a natural endosomal environment, in particular the likely presence of mRNA escape into the solution followed by rebinding of monomeric mRNA to the then ionizable-lipid-containing endosomal membrane, as well as the presence of multiple entangled mRNA molecules that remain associated with the site of fusion (Figure 3). This opens up for two plausible mRNA escape mechanisms: i) disentanglement from mRNA at the site of fusion into the neutral cytosolic environment, or ii) escape of individual mRNA bound to the endsomal membrane through pores in the membrane caused by endosomal remodeling and disintegration caused by transfer of ionized lipids from the LNP.

It is in this context worth noting that in our experiments, it seems that only a minor fraction of the mRNA that escapes into the solution upon LNP fusion (Figure 3) rebinds to endosomal membrane mimic. However, given the low surface-to-liquid volume ratio in the microfluidic channel, this is expected. If this process occurs for LNPs bound to the membrane of a closed endosomal compartment with a sub-micrometer dimension, all suspended mRNA molecules would reside in close proximity to the inner endosomal membrane with high probability to bind, mediated through for example ionized lipids that escape into the membrane during LNP fusion or because suspended mRNA is complexed with ionized lipids. Considering the presence of this phenomenon, it is not obvious whether mRNA translocation into the neutral cytosolic environment, which is expected to be a very rare process, is more likely to occur through translocation of individual endosomal-membrane-bound mRNA in response to endosomal damage,^17,21^ or whether mRNA manages to escape prior to endosomal damage from the entangled mRNA state that we observed remaining at the site of fusion (Figure 2). Further, the fact that the complex salt formed between mRNA and DLin-MC3-DMA at acidic conditions is not instantaneously reversed at neutral pH^48^ is yet another factor that may contribute to the low transfection efficiency, since it may hamper the efficiency of the ribosomal protein synthesis or lead to undesired mRNA association to internal cellular membrane compartments.

We also note that there is a wide distribution in the wait-time between a reduction in pH and onset of LNP fusion, lasting between tens to hundreds of seconds depending on pH, size and type of LNP. Further, around 40% of the most fusogenic type of LNP (low-DSPC LNPs) do not undergo fusion even when the pH was eventually reduced to 5.6 (Figures 1 and 4), at which the majority of the ionizable lipids are expected to be ionized. Thus, even if ionizable-lipid containing LNPs serve to provide very efficient mRNA encapsulation, typically exceeding 95%, and the correct choice of helper lipids can provide sufficient stability and circulation times for efficient cellular uptake, we dare to conclude from this study that there are significant variations between individual LNPs with respect to their fusion capacity. In addition, the fate of individual mRNA molecules encapsulated in the very same LNP may differ. This, in turn, calls for efforts devoted to refined LNP fabrication protocols with the capacity to produce LNP batches with significantly reduced variations. Further, in compliance with cellular data, almost one order of magnitude more efficient fusion was observed for LNPs with reduced surface density of gel-phase forming DSPC lipids, despite that both low- and high DSPC LNPs display identical charge titration curves using the TNS assay. This suggests that assays of the type reported in this work, that goes beyond characterization of LNP size, encapsulation efficiency, structure and surface charge could serve as a helpful tool not only to provide mechanistic insights regarding the physicochemical mechanisms that underpin pH-induced LNP fusion with anionic membranes, but also to identify formulations with preferred features and the desired functional response.

## MATERIALS AND METHODS

### LNP composition

The low- and high-DSPC LNP formulations contained ionizable cationic lipid O-(Z,Z,Z,Z-heptatriaconta-6,9,26,29-tetraem-19-yl)-4-(N,N-dimethylamino)butanoate (DLin-MC3-DMA), 1,2-distearoyl-sn-glycero-3-phosphocholine (DSPC), cholesterol, 1,2-dimyristoyl-sn-glycero-3-phosphoethanolamine-N-[methoxy(polyethyleneglycol)-2000] (DMPE-PEG2000), 1,2-distearoyl-sn-glycero-3-phosphoethanolamine-N-[biotinyl(polyethylene glycol)-2000] (DSPE-PEG(2000) Biotin) and 1,2-dioleoyl-sn-glycero-3-phosphoethanolamine-N-(lissamine rhodamine B sulfonyl) (Rhod-DOPE) in molar ratios given in Table S1. The LNP cargo contained Cy5-labeled (TriLink Bio Technologies) and non-labeled eGFP-encoding mRNA at a 1:4 molar ratio. According to the manufacturer the ratio of Cy5-labeled and unlabeled uridine (U) is 1:3. The open reading frame of eGFP contains 103 U which gives 26 Cy5-U. Additionally, there are another 277 nucleotides in the full sequence with ∼120-150 estimated to make up the poly(A)-tail. ^52^ If one assumes that 25% of the remaining 120-160 nucleotides are U we expect an additional 7-9 Cy5-U, resulting in a total of ∼ 34 Cy5-U per mRNA. The PolyA- and calcein-containing LNPs were made from aqueous solution containing 30 mM calcein and with eGFP-mRNA replaced with PolyA (Table S1).

### LNP Preparation and Characterization

Low- and high DSPC LNPs were prepared using the NanoAssemblr Benchtop device, while PolyA- and calcein-containg LNPs were made using the NanoAssemblr® Spark™ device (both from Precision Nanosystems Inc, Canada). Briefly, stocks of lipids were dissolved in ethanol and mixed in the appropriate molar ratios to obtain a lipid concentration of 12.5 mM. mRNA or PolyA were diluted in RNase free citrate buffer (TekNova) of 50 mM at pH 3.0 to obtain a mRNA/PolyA:lipid weight ratio of 10:1. The aqueous and ethanol solutions were mixed in a 3:1 volume ratio through a microfluidic cartridge of the Benchtop device at a flow rate of 12 mL min^-1^. LNPs were dialyzed overnight against 600× sample volume nuclease-free PBS using Slide-A-Lyzer G2 dialysis cassettes (Thermo Scientific) with a molecular weight cutoff of 10 K. The collected LNPs with a mRNA concentration of 0.1 mg/mL were filtered through a sterile filter (0.2 µm) prior to use. The size of LNPs was determined by DLS measurements using a ZetaSizer Nano ZS from Malvern Instruments Ltd. The encapsulation efficiency was determined using the RiboGreen assay (ThermoFisher). LNP size and concentration also determined using nanoparticle tracking analysis (NTA) using a Nanosight LM10 device with a Hamamatsu C11440-50B/A11893-02 camera. The anionic fluorescent dye 2-(p-toluidino)-6-napthalene sulfonic acid (TNS) measurements were performed in a 384-well format with a buffer containing 20 mM phosphate tribasic, 25 mM ammonium citrate, 20 mM ammonium acetate and 150 mM sodium chloride, with a pH ranging from 2 to 11. The molar ratio of total lipid:TNS dye was kept at 4.25 and the total lipid concentration in each well was kept at 7.3 µM. All measurements were performed at room temperature within 10 minutes of preparation using a fluorescence plate reader (BMG Labtech) with excitation at 340 nm and emission at 460 nm.

### Endosomal membrane mimic composition

Lipids used to produce the lipid vesicles used for SLB formation were purchased from Avanti® Polar Lipids, Inc. The lipid vesicles were produced using the lipid film hydration and extrusion method. In brief, 1-palmitoyl-2-oleoyl-sn-glycero-3-phosphocholine (POPC), 1-palmitoyl-2-oleoyl-sn-glycero-3-phospho-L-serine (POPS), 1,2-dioleoyl-sn-glycero-3-phosphoethanolamine-N-(cap biotinyl) (Biotin-Cap-DOPE) and 1,2-dioleoyl-sn-glycero-3-phosphoethanolamine-N-(7-nitro-2-1,3-benzoxadiazol-4-yl) (NBD-DOPE), suspended in chloroform at concentration of 10, 10, 0.5 and 1 mg mL^-1^, respectively. 186.30 µL of POPC (93.45 mol%), 12.34 µl of POPS (6 mol%), 2.9 µL of Biotin-Cap-DOPE(0.05 mol%), and 12.12 µL of NBD-DOPE(0.5 mol%) were mixed and dried in vacuum overnight. The lipid film was rehydrated with PBS for 1 hour to a total lipid concentration of 2 mg mL^-1^. The solution was subsequently extruded 21 times using a mini extruder (Avanti Lipids Inc., Alabaster, AL, USA) with 50 and 30 nm polycarbonate membranes (Whatman, Maidstone, UK) to form the vesicles with the size of approximately 100 nm. The vesicle solution was stored at 4 °C for later use. The zeta potential ζ for the lipid vesicles was determined using ZetaSizer (Malvern) to -22.6 ± 2.05 V.

### Nanoporous silica thin film formation

Nanoporous silica thin films were synthesized following a modified method of Alberius et al. ^53^ Briefly, 0.28 g of Poly(ethylene glycol)-block-poly(propylene glycol)-block-poly(ethylene glycol) (P123, Sigma-Aldrich) was dissolved in 1.33 g of ethanol (99.5%, Solveco) in a glass vial. This mixture was stirred using a magnetic stirrer at room temperature until completely dissolved. In a separate vial, 1.73 g of Tetraethylorthosilicate (TEOS, 98%, Sigma-Aldrich) and 2 g of ethanol were combined and stirred at 300 rpm with a magnetic stirrer. Subsequently, 0.9 g of 0.01 M HCl (Sigma-Aldrich) was added dropwise to this mixture and stirred continuously for 20 minutes. After this period, the P123 solution was mixed with theTEOS solution. This silica precursor solution was then stirred at room temperature at 300 rpm for 20 minutes to achieve a homogeneous and clear solution.

The silica precursor solution was deposited onto borosilicate cover glasses (Menzel-Gläser, D263®, number 1) through spin-coating at 4000 rpm (WS-650, Laurell Technologies Corporation). This was done immediately after submerging the glasses in an ETOH-NaOH (5:1) cleaning solution for 5 minutes, followed by a thorough rinse with ultrapure water (MilliQ, Merck Millipore) and drying using nitrogen gas. The coated glasses were then left in the dark to age at room temperature for 24 hours. The templating agent was removed by gradual heating at a rate of 1 °C per minute from room temperature to 400 °C, and maintaining this temperature for 4 hours before allowing cooling to room temperature. Top view SEM (Scanning Electron Microscopy) analysis of the silica thin films formed on silicon wafers was performed using a Leo Ultra 55 FEG SEM (Zeiss) at an operating voltage of 1 kV.

### Endosomal model membrane formation

To prepare the microfluidic channel, the Ibidi sticky microfluidic channels (3.8×17×0.4 mm in width × length × height, Ibidi® cell in focus, Gräfelfing, Germany) were attached to the nanoporous substrate after first being cleaned thoroughly by ethanol and water followed by two subsequent steps of UV ozone treatment (for ∼20 minutes), MilliQ/ethanol rinsing, and nitrogen drying. Anionic SLBs were formed on the porous substrate by injecting the lipid vesicle suspension (diluted to a lipid concentration of 200 µg mL^-1^) in PBS. NeutrAvidin suspended in PBS (∼20 µg mL^-1^) was injected in the channel at a flow rate of 50 µl per minute for 10 minutes followed by thorough rinsing for 5 minutes with citrate-phosphate buffer (pH 7.5). The LNP stock solution was diluted 1000 times in phosphate-citrate buffer (pH 7.5) and injected at a flow rate of 140 µL min^-1^ until a suitable LNP coverage was obtained (see main text), followed by 5 minutes rinsing in pure buffer. To vary the pH a citrate-phosphate buffer was used to vary the pH from 7.5 to 5.6.

### Fluorescence microscopy

For time resolved imaging, the microfluidic system was mounted on an inverted Eclipse Ti-E microscope (Nikon Corporation, Minato City, Japan) equipped with a CFI Apo TIRF 60× (NA: 1.49) oil immersion objective (Nikon Corporation, Tokyo, Japan). A FITC filter set (Semrock, Sandwich, IL, USA) was used to excite the NBD-DOPE dyes for visualizing lipid vesicle adsorption and SLB formation on the nanoporous silica substrate. In addition, the continuity and fluidity of the bilayer was evaluated using FRAP assessment by bleaching NBD lipids in a circular region (spot) of the SLB with a solid-state light source (Lumencor Spectra X-LED) at a wavelength of 531 nm followed by imaging of the fluorescence recovery with 12 fps. FRAP data was analyzed using a custom written code ^35^ in MATLAB (MathWorks. Inc., USA). LNP binding to the anionic SLB was using TIRF microscopy utilizing a TRITC filter set (Semrock, Sandwich, IL, USA) for the Rhod-DOPE dyes (excitation:565nm; emission:590 nm) and a Cy5 ET filter set (F46-006 ET-set, Chroma Technology Corporation, USA) for the Cy5 dyes (excitation: 640nm; emission: 670 nm), conjugated to the mRNA cargo. Imaging was performed for Rhod-DOPE and Cy5-mRNA separately, at an exposure time of 50 ms with a frame rate of 3 fps and 2 fps, respectively. Simultaneous imaging of Rhod-DOPE and Cy5-mRNA was performed using an image splitter (OptoSplit II, Cairn research) at 20 fps.

### Image Analysis

The positions of individual signals in the fluorescence micrographs were determined using a threshold-based maxima detection, followed by a sub-pixel position determination employing radial symmetry characteristics.^54^ For time-resolved videos, where intensity extraction for numerous signals was done (as Figure 1, 3, and 4), the signal positions were linked into trajectories using the Hungarian algorithm ^55^. The emission intensities were extracted from the background subtracted micrographs as the sum of pixel values in a quadratic area with the center defined by the position, and the side length selected to reflect the average extension of signals in the micrographs.

Fusion events were detected automatically using a custom-written script, with a fusion event defined as a relative intensity change of >30% within a <5 consecutive frames, generally also coinciding with loss of the particle trajectory. For data displaying particularly low fusion efficiency, e.g., at pH decrease from 7.5 to 6.6, fusion events were marked manually using the Point Picker plugin (Philippe Thévenaz, Biomedical Imaging Group, Swiss Federal Institute of Technology, Lausanne) of ImageJ and later matched with specific particle trajectories.

The temporal evolution of intensity profiles used for extraction of lateral Rhod-DOPE diffusivity *D*, a 2-dimensional Gaussian function was fitted to a 34×34 pixels centered around the intensity profile maximum. The emission intensity of the particle and the region surrounding the particle were extracted as the sum of values of the Gaussian fit in an area defined by a maximum distance of 4 pixels, and a distance between 4 and 16 pixels to the signal center, respectively. The emission intensity of the particle and the surrounding area both represent background-subtracted intensities as the Gaussian function was shifted to zero offset prior to a fusion event. The relative change of emission intensity upon fusion was determined from the change in total emission intensity, obtained from the integral of the Gaussian fit. The lateral diffusion was determined as previously described ^56^ by extracting the orthogonal position variations Δx and Δy, subsequently used to estimate the mean square displacement MSDi=< Δi>^2^, from which the 1D diffusion coefficient was obtained as *D*_i_ = MSD_i_/(4 Δ*t*), for i=x,y. Finally, the 2D diffusion coefficient *D* was calculated as arithmetic average of *D*_x_ and *D*_y_. All data analysis was performed using MATLAB (MathWorks. Inc., USA).

## Acknowledgements

Authors would like to acknowledge the Swedish Foundation of Strategic Research for financing the project and all the members of Industrial Research Centre “FoRmulaEx” (IRC15-0065), The Swedish Research Council (#2022-05016) and the Wallenberg Foundation (#2019-0577) for financial support. We would like to thank Simon Isaksson for assistance with the nanoporous silica formation protocols and Mokhtar Mapar for valuable discussions.

## References

1 Damase, T. R. et al. The Limitless Future of RNA Therapeutics. Front. Bioeng. Biotechnol. 9, 24 (2021). 10.3389/fbioe.2021.628137

2 Paunovska, K., Loughrey, D. & Dahlman, J. E. Drug delivery systems for RNA therapeutics. Nat Rev Genet 23, 265–280 (2022). 10.1038/s41576-021-00439-4

3 Hajj, K. A. & Whitehead, K. A. Tools for translation: non-viral materials for therapeutic mRNA delivery. Nat Rev Mater 2 (2017). 10.1038/natrevmats.2017.56

4 Phua, K. K., Leong, K. W. & Nair, S. K. Transfection efficiency and transgene expression kinetics of mRNA delivered in naked and nanoparticle format. J Control Release 166, 227–233 (2013). 10.1016/j.jconrel.2012.12.029

5 Yin, H. et al. Non-viral vectors for gene-based therapy. Nat Rev Genet 15, 541–555 (2014). 10.1038/nrg3763

6 Lagache, T., Danos, O. & Holcman, D. Modeling the step of endosomal escape during cell infection by a nonenveloped virus. Biophys J 102, 980–989 (2012). 10.1016/j.bpj.2011.12.037

7 Nayak, S. & Herzog, R. W. Progress and prospects: immune responses to viral vectors. Gene Ther 17, 295–304 (2010). 10.1038/gt.2009.148

8 Schott, J. W., Morgan, M., Galla, M. & Schambach, A. Viral and Synthetic RNA Vector Technologies and Applications. Mol Ther 24, 1513–1527 (2016). 10.1038/mt.2016.143

9 Thomas, S. J. et al. Safety and Efficacy of the BNT162b2 mRNA Covid-19 Vaccine through 6 Months. N Engl J Med 385, 1761–1773 (2021). 10.1056/NEJMoa2110345

10 Widge, A. T. et al. Durability of Responses after SARS-CoV-2 mRNA-1273 Vaccination. N Engl J Med 384, 80–82 (2021). 10.1056/NEJMc2032195

11 Cullis, P. R. & Hope, M. J. Lipid Nanoparticle Systems for Enabling Gene Therapies. Mol Ther 25, 1467–1475 (2017). 10.1016/j.ymthe.2017.03.013

12 Cheng, X. & Lee, R. J. The role of helper lipids in lipid nanoparticles (LNPs) designed for oligonucleotide delivery. Adv Drug Deliv Rev 99, 129–137 (2016). 10.1016/j.addr.2016.01.022

13 Kulkarni, J. A., Cullis, P. R. & van der Meel, R. Lipid Nanoparticles Enabling Gene Therapies: From Concepts to Clinical Utility. Nucleic Acid Ther 28, 146–157 (2018). 10.1089/nat.2018.0721

14 Leung, A. K., Tam, Y. Y., Chen, S., Hafez, I. M. & Cullis, P. R. Microfluidic Mixing: A General Method for Encapsulating Macromolecules in Lipid Nanoparticle Systems. J Phys Chem B 119, 8698–8706 (2015). 10.1021/acs.jpcb.5b02891

15 Truong, L. B., Medina-Cruz, D. & Mostafavi, E. Current state of RNA delivery using lipid nanoparticles to extrahepatic tissues: A review towards clinical translation. Int. J. Biol. Macromol. 242, 8 (2023). 10.1016/j.ijbiomac.2023.125185

16 Dowdy, S. F., Setten, R. L., Cui, X. S. & Jadhav, S. G. Delivery of RNA Therapeutics: The Great Endosomal Escape! Nucleic Acid Ther 32, 361–368 (2022). 10.1089/nat.2022.0004

17 Wittrup, A. et al. Visualizing lipid-formulated siRNA release from endosomes and target gene knockdown. Nature Biotechnology 33, 870-+ (2015). 10.1038/nbt.3298

18 Mellman, I., Fuchs, R. & Helenius, A. Acidification of the Endocytic and Exocytic Pathways. Annu Rev Biochem 55, 663–700 (1986). 10.1146/annurev.biochem.55.1.663

19 Falguieres, T., Luyet, P. P. & Gruenberg, J. Molecular assemblies and membrane domains in multivesicular endosome dynamics. Exp Cell Res 315, 1567–1573 (2009). 10.1016/j.yexcr.2008.12.006

20 Akinc, A. et al. The Onpattro story and the clinical translation of nanomedicines containing nucleic acid-based drugs. Nat Nanotechnol 14, 1084–1087 (2019). 10.1038/s41565-019-0591-y

21 Munson, M. J. et al. A high-throughput Galectin-9 imaging assay for quantifying nanoparticle uptake, endosomal escape and functional RNA delivery. Commun Biol 4, 211 (2021). 10.1038/s42003-021-01728-8

22 Gilleron, J. et al. Image-based analysis of lipid nanoparticle-mediated siRNA delivery, intracellular trafficking and endosomal escape. Nat Biotechnol 31, 638–646 (2013). 10.1038/nbt.2612

23 Kauffman, K. J. et al. Optimization of Lipid Nanoparticle Formulations for mRNA Delivery in Vivo with Fractional Factorial and Definitive Screening Designs. Nano Lett. 15, 7300–7306 (2015). 10.1021/acs.nanolett.5b02497

24 Spadea, A. et al. Nucleic Acid-Loaded Lipid Nanoparticle Interactions with Model Endosomal Membranes. ACS Appl Mater Interfaces 14, 30371–30384 (2022). 10.1021/acsami.2c06065

25 Aliakbarinodehi, N. et al. Interaction Kinetics of Individual mRNA-Containing Lipid Nanoparticles with an Endosomal Membrane Mimic: Dependence on pH, Protein Corona Formation, and Lipoprotein Depletion. ACS Nano 16, 20163–20173 (2022). 10.1021/acsnano.2c04829

26 Flavier, K. M. & Boxer, S. G. Vesicle Fusion Mediated by Solanesol-Anchored DNA. Biophys J 113, 1260–1268 (2017). 10.1016/j.bpj.2017.05.034

27 Karatekin, E. et al. A fast, single-vesicle fusion assay mimics physiological SNARE requirements. Proc Natl Acad Sci U S A 107, 3517–3521 (2010). 10.1073/pnas.0914723107

28 Claesson, M., Frost, R., Svedhem, S. & Andersson, M. Pore spanning lipid bilayers on mesoporous silica having varying pore size. Langmuir 27, 8974–8982 (2011). 10.1021/la201411b

29 Raman, N. K., Anderson, M. T. & Brinker, C. J. Template-based approaches to the preparation of amorphous, nanoporous silicas. Chem Mater 8, 1682–1701 (1996). DOI 10.1021/cm960138+

30 Joyce, P. et al. TIRF Microscopy-Based Monitoring of Drug Permeation Across a Lipid Membrane Supported on Mesoporous Silica. Angew Chem Int Ed Engl 60, 2069–2073 (2021). 10.1002/anie.202011931

31 Chen, D., Ganesh, S., Wang, W. & Amiji, M. The role of surface chemistry in serum protein corona-mediated cellular delivery and gene silencing with lipid nanoparticles. Nanoscale 11, 8760–8775 (2019). 10.1039/c8nr09855g

32 Akinc, A. et al. Targeted delivery of RNAi therapeutics with endogenous and exogenous ligand-based mechanisms. Mol Ther 18, 1357–1364 (2010). 10.1038/mt.2010.85

33 Yanez Arteta, M., et al. Successful reprogramming of cellular protein production through mRNA delivered by functionalized lipid nanoparticles. Proc Natl Acad Sci U S A 115, E3351–E3360 (2018). 10.1073/pnas.1720542115

34 Kobayashi, T. et al. Separation and characterization of late endosomal membrane domains. J Biol Chem 277, 32157–32164 (2002). 10.1074/jbc.M202838200

35 Jonsson, P., Jonsson, M. P., Tegenfeldt, J. O. & Hook, F. A method improving the accuracy of fluorescence recovery after photobleaching analysis. Biophys J 95, 5334–5348 (2008). 10.1529/biophysj.108.134874

36 Eastman, S. J., Hope, M. J. & Cullis, P. R. Transbilayer Transport of Phosphatidic-Acid in Response to Transmembrane Ph Gradients. Biochemistry-Us 30, 1740–1745 (1991). 10.1021/bi00221a002

37 Jayaraman, M. et al. Maximizing the Potency of siRNA Lipid Nanoparticles for Hepatic Gene Silencing In Vivo. Angew Chem Int Edit 51, 8529–8533 (2012). 10.1002/anie.201203263

38 Axelrod, D., Burghardt, T. P. & Thompson, N. L. Total internal reflection fluorescence. Annu Rev Biophys Bioeng 13, 247–268 (1984). 10.1146/annurev.bb.13.060184.001335

39 Zhang, Y., Li, Q., Dong, M. & Han, X. Effect of cholesterol on the fluidity of supported lipid bilayers. Colloids Surf B Biointerfaces 196, 111353 (2020). 10.1016/j.colsurfb.2020.111353

40 Chen, C., Corry, B., Huang, L. & Hildebrandt, N. FRET-Modulated Multihybrid Nanoparticles for Brightness-Equalized Single-Wavelength Barcoding. J Am Chem Soc 141, 11123–11141 (2019). 10.1021/jacs.9b03383

41 Hannestad, J. K. et al. Single-vesicle imaging reveals lipid-selective and stepwise membrane disruption by monomeric alpha-synuclein. P Natl Acad Sci USA 117, 14178–14186 (2020). 10.1073/pnas.1914670117

42 Sjoberg, M. et al. Time-Resolved and Label-Free Evanescent Light-Scattering Microscopy for Mass Quantification of Protein Binding to Single Lipid Vesicles. Nano Lett. 21, 4622–4628 (2021). 10.1021/acs.nanolett.1c00644

43 Li, M., Harbron, R. L., Weaver, J. V., Binks, B. P. & Mann, S. Electrostatically gated membrane permeability in inorganic protocells. Nat Chem 5, 529–536 (2013). 10.1038/nchem.1644

44 Zhang, Y. et al. The development of an in vitro assay to screen lipid based nanoparticles for siRNA delivery. J Control Release 174, 7–14 (2014). 10.1016/j.jconrel.2013.11.006

45 Kho, K. W., Berselli, G. B. & Keyes, T. E. A Nanoplasmonic Assay of Oligonucleotide-Cargo Delivery from Cationic Lipoplexes. Small 17 (2021). 10.1002/smll.202008155

46 Kamanzi, A. et al. Simultaneous, Single-Particle Measurements of Size and Loading Give Insights into the Structure of Drug-Delivery Nanoparticles. ACS Nano 15, 19244–19255 (2021). 10.1021/acsnano.1c04862

47 Tsui, F. C., Ojcius, D. M. & Hubbell, W. L. The Intrinsic Pka Values for Phosphatidylserine and Phosphatidylethanolamine in Phosphatidylcholine Host Bilayers. Biophysical Journal 49, 459–468 (1986). 10.1016/S0006-3495(86)83655-4

48 Philipp, J. et al. pH-dependent structural transitions in cationic ionizable lipid mesophases are critical for lipid nanoparticle function. Proc Natl Acad Sci U S A 120, e2310491120 (2023). 10.1073/pnas.2310491120

49 Li, M. et al. Brush Conformation of Polyethylene Glycol Determines the Stealth Effect of Nanocarriers in the Low Protein Adsorption Regime. Nano Lett 21, 1591–1598 (2021). 10.1021/acs.nanolett.0c03756

50 Kolter, T. & Sandhoff, K. Lysosomal degradation of membrane lipids. Febs Lett 584, 1700–1712 (2010). 10.1016/j.febslet.2009.10.021

51 Francia, V., Schiffelers, R. M., Cullis, P. R. & Witzigmann, D. The Biomolecular Corona of Lipid Nanoparticles for Gene Therapy. Bioconjugate Chem 31, 2046–2059 (2020). 10.1021/acs.bioconjchem.0c00366

52 Sahin, U., Kariko, K. & Tureci, O. mRNA-based therapeutics--developing a new class of drugs. Nat Rev Drug Discov 13, 759–780 (2014). 10.1038/nrd4278

53 Alberius, P. C. A. et al. General predictive syntheses of cubic, hexagonal, and lamellar silica and titania mesostructured thin films. Chem Mater 14, 3284–3294 (2002). 10.1021/cm011209u

54 Parthasarathy, R. Rapid, accurate particle tracking by calculation of radial symmetry centers. Nat Methods 9, 724–726 (2012). 10.1038/nmeth.2071

55 Kuhn, H. W. The Hungarian Method for the assignment problem. Nav Res Log 52, 7–21 (2005). 10.1002/nav.20053

56 Block, S., Fast, B. J., Lundgren, A., Zhdanov, V. P. & Hook, F. Two-dimensional flow nanometry of biological nanoparticles for accurate determination of their size and emission intensity. Nat Commun 7, 12956 (2016). 10.1038/ncomms12956

