## Supporting Information Figures for "Time Resolved Inspection of Ionizable-Lipid Facilitated Lipid Nanoparticle Disintegration and Cargo Release at an Endosomal Membrane Mimic"

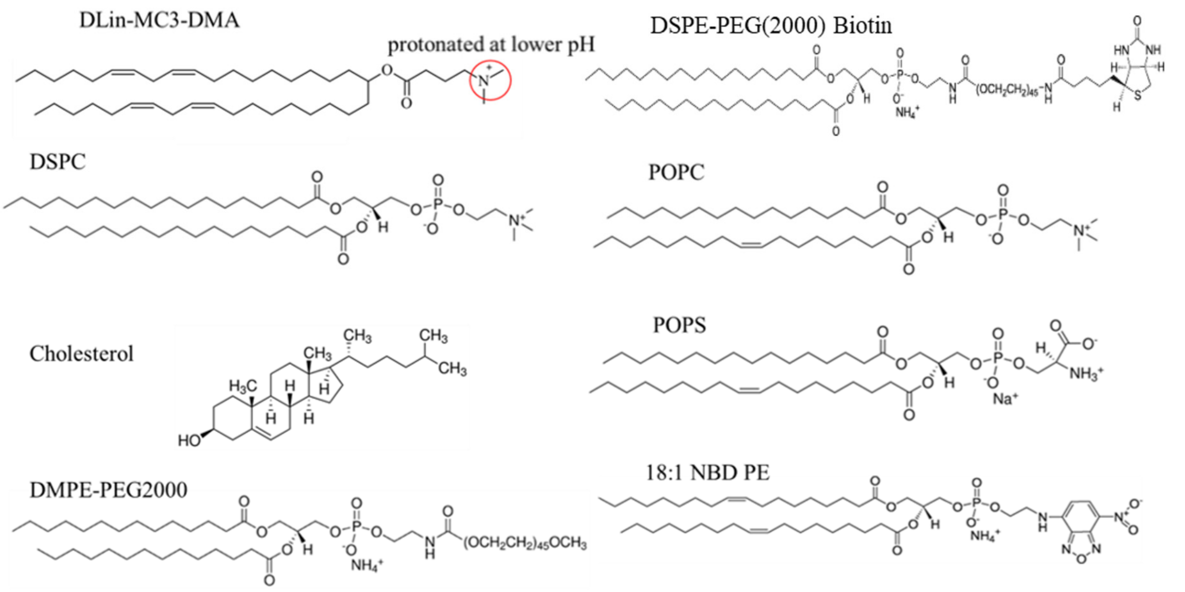


**Figure S1**. Structures of lipids used in LNP preparations.

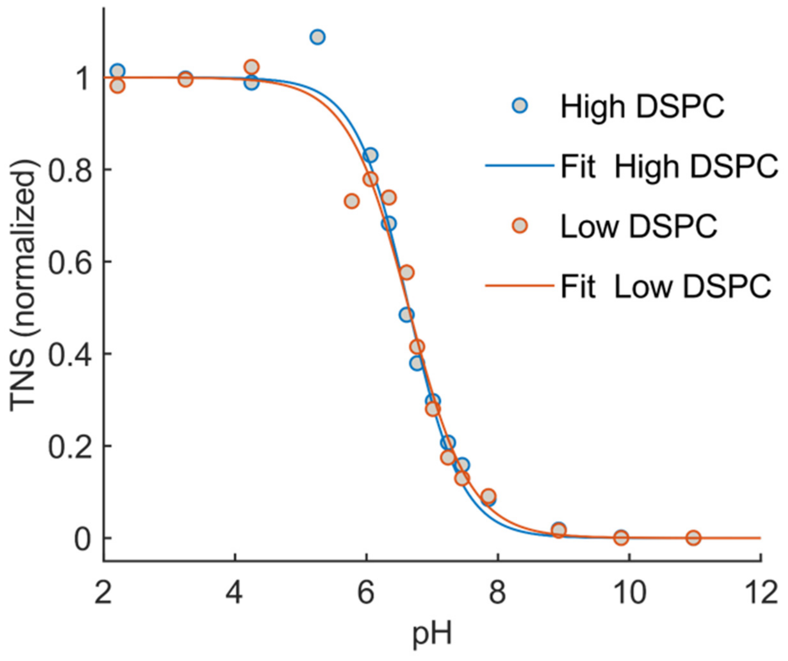


**Figure S2** In situ TNS fluorescence titration of low- (blue circles) and high-DSPC (red circles) LNPs. Duplicate measurements were averaged and fitted (solid lines) to a three-parameter sigmoidal.

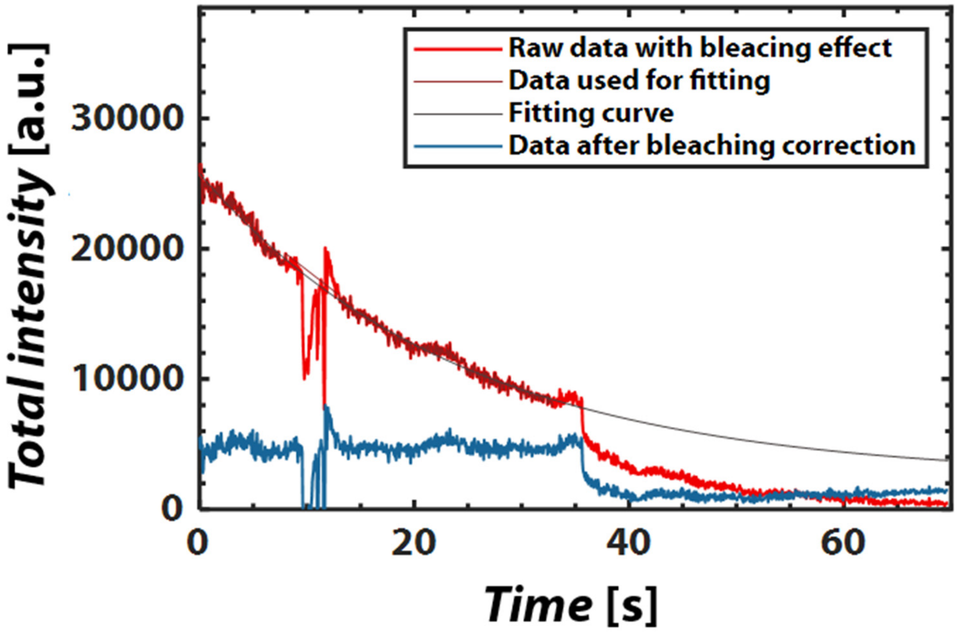


**Figure S3**. Representative example of time-resolved total Cy5-mRNA emission intensity changes upon reduction of the pH from 6.0 to 5.6 extracted from a single LNP visualized in Epi mode. The red curve represents raw data, and the blue curve represents the bleaching-corrected data.

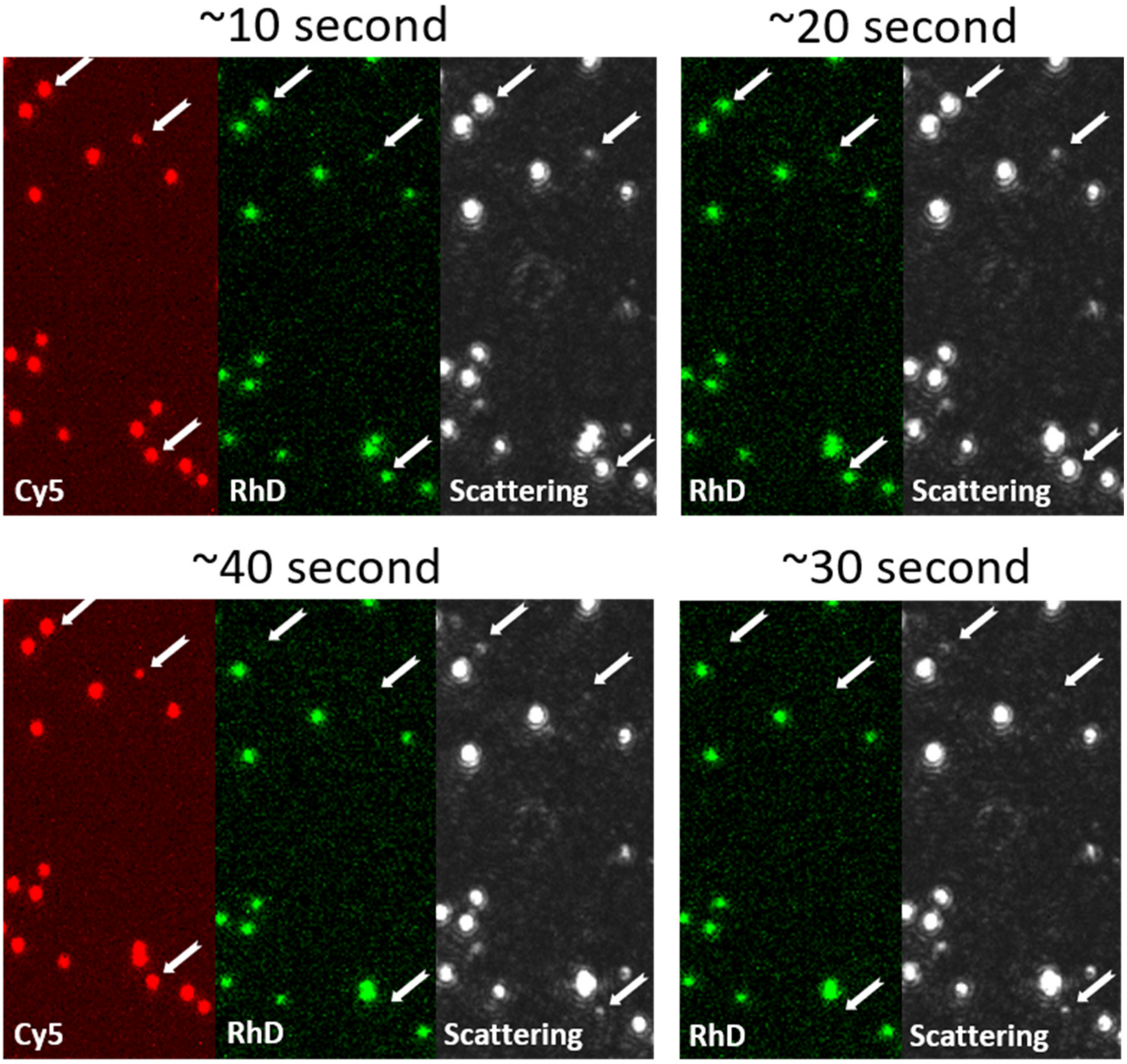


**Figure S4.** Time resolved dual-mode fluorescence and label-free-scattering micrographs measured upon a reduction of the pH from 6.0 to 5.6 for low-DSPC LNPs tethered via NeuraAvidin to the same type anionic SLB as used in the main text, but here formed on a planar glass of a Nanolyze Sense waveguide chip (Nanolyze AB)[1]. An Olympus BX61 microscope was used for image acquisition (objective: 60X, NA 1.0, camera: ORCA-Flash 4.0 V2.0 CMOS, image splitter: Hamamatsu W-VIEW GEMINI, lasers: Cobolt 06-MLD, $\lambda$ = 488 nm, and OZ 3000, $\lambda$ = 635 nm). Both the DOPE-Rhod and the scattering signal display a significant decrease at ~30s, consistent with fusion, while the reduction in the Cy5-mRNA signal is less dramatic. The magnitude of the decrease in the scattering signal is >90%.

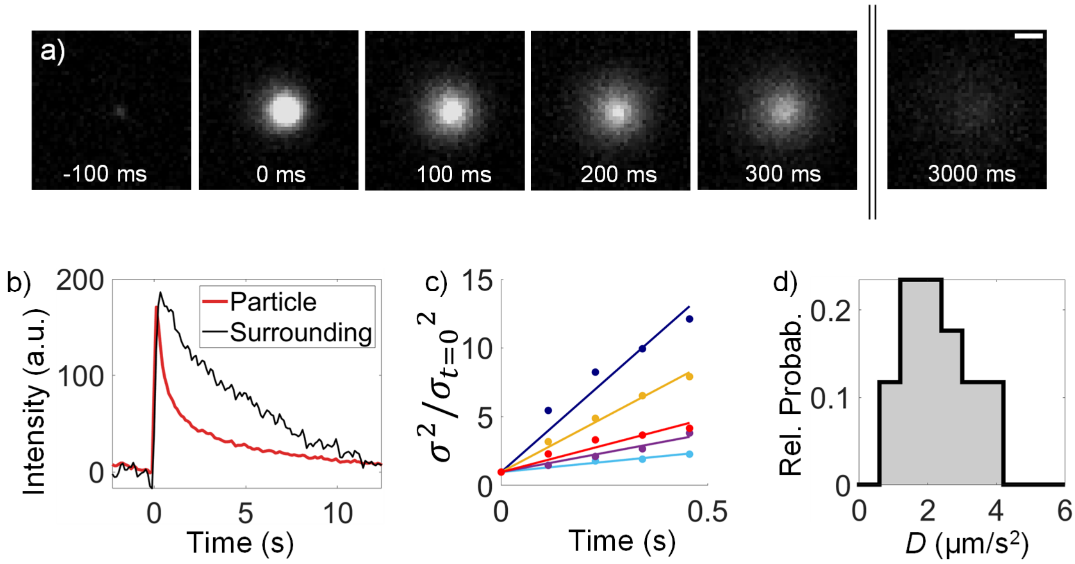


**Figure S5.** TIRF micrographs of a tethered calcein-containing LNPs, c.f. Fig. 2 in the main text (scalebar 1 µm). **b)** The time-evolution of the total intensity emission (solid lines) and the emission from an area surrounding the LNP docking site (dashed lines) extracted from background subtracted emission profiles represented by two-dimensional Gaussian profiles (see Fig. 2 in main text). The initial increase in the total intensity is attributed to calcein dequenching upon LNP collapse, followed by lateral escape of calcein-DLin-MC3-DMA complexes in the endosomal membrane mimic. The higher total intensity in the area surrounding the LNP is attributed to a combination of a temporal increase due to dequenching and higher illumination intensity experienced by the fluorophore as it moves closer to the glass interface upon LNP collapse. **c)** $\sigma^2/\sigma_{t=0}^2$ versus time, with the variance obtained from the Gaussian representation of the LNPs **d)** Distribution of diffusion constants, *D,* obtained from 17 different LNP-variances, $\sigma^2 = 2Dt$, from c).

**Table S1**. Lipid compositions and characteristics of all LNP formulations used in the presented study. mRNA encapsulation and concentration were determined using the RiboGreen assay. Size and concentration characterization were conducted using dynamic light scattering (DLS) and nanoparticle tracking analysis (NTA).

| | low-DSPC LNPs | high-DSPC LNPs | calcein LNPs |
|---|---|---|---|
| Composition (lipids, mol%) | | | |
| DLin-MC3-DMA | 53.47 | 50 | 53.47 |
| DSPC | 4.65 | 10 | 4.65 |
| Chol | 41.114 | 39.684 | 41.174 |
| DSPE-PEG(2000) Biotin | 0.006 | 0.006 | 0.006 |
| DMPE-PEG(2000) | 0.7 | 0.25 | 0.7 |
| Rhod-DOPE | 0.06 | 0.06 | NA |
| Characteristics | | | |
| mRNA encapsulation (%) | 97 | 98 | 34** |
| mRNA concentration (μg $mL^{-1}$) | 0.053 | 0.051 | 0.17 |
| LNP diameter (nm) | 139 | 142 | 145.5 |
| LNP concentration (particles $mL^{-1}$)* | $1.26\times10^{12}$ | $0.82\times10^{12}$ | $2.20\times10^{12}$ |
| PDI | 0.031 | 0.019 | <0.5* |

*Measured by NTA

** PolyA

**References**


1. Agnarsson, B. *et al.* Low-temperature fabrication and characterization of a symmetric hybrid organic-inorganic slab waveguide for evanescent light microscopy. *Nano Futures* **2**, (2018).
